# A subset of lung cancer cases shows robust signs of homologous recombination deficiency associated genomic mutational signatures

**DOI:** 10.1101/576223

**Authors:** Miklos Diossy, Zsofia Sztupinszki, Judit Borcsok, Marcin Krzystanek, Viktoria Tisza, Sandor Spisak, Orsolya Rusz, Jozsef Timar, István Csabai, Judit Moldvay, Anders Gorm Pedersen, David Szuts, Zoltan Szallasi

## Abstract

**Background:** Consistent with their assumed mechanism of action, PARP inhibitors show significant therapeutic efficacy in breast, ovarian and prostate cancer, which are the solid tumor types most often associated with the loss of function of key homologous recombination genes. It remains unknown, however, how frequent homologous recombination deficiency (HRD) is in other solid tumor types. Since it is well established, that HRD induces specific DNA aberration profiles and genomic scars that can be captured by various next-generation sequencing (NGS) based biomarkers, it is possible to assess the presence or absence of this DNA repair pathway aberration in any given tumor biopsy.

**Methods:** We derived the various HRD associated mutational signatures from whole genome and whole exome sequencing data in the lung adenocarcinoma (LUAD) and lung squamous carcinoma (LUSC) cases from TCGA, in a patient of ours with stage IVA lung cancer with exceptionally good response to platinum-based therapy and in lung cancer cell lines.

**Results:** We have found evidence that a subset of the investigated cases shows robust signs of HR deficiency, some of which exhibiting similar patterns to those with a complete loss of function of either BRCA1 or BRCA2 genes, however, without any signs of genetic alterations being present in either of those genes. The extreme platinum responder case also showed a robust HRD associated genomic mutational profile. HRD associated mutational signatures were also associated with PARP inhibitor sensitivity in lung cancer cell lines.

**Conclusions:** Lung cancer cases with high levels of HRD associated mutational signatures could be candidates for PARP inhibitor treatment, and in general, the prioritization of patients for clinical trials might be achieved using the combined analysis of the HRD-related next-generation sequencing-based mutational signatures.

## Background

PARP inhibitors are a new class of cancer therapeutic agents that are most effective in tumors lacking the homologous recombination mediated DNA repair pathway[1]. They are approved either in BRCA1/2 mutant tumors or in solid tumor types most often associated with lack of HR in the form of BRCA1/2 function deficiency[1]. It is possible, however, that other tumor types not associated with germline BRCA1/2 mutations may also be HR deficient. Non-small cell lung cancers, for example, show somatic mutations in the BRCA1/2 genes in 5-10% of the cases[2], and they also harbor mutations in various DNA damage checkpoint genes [3, 4]. Recently, it was estimated that 2-3 % of NSCLC cases in fact harbor somatic or germline loss of function mutations of the BRCA1/2 genes and those are often associated with loss of heterozygosity as well [5]. We were seeking further evidence that such tumor cases are in fact homologues recombination deficient by investigating the next generation sequencing-based DNA aberrations profiles of TCGA cohorts.

Loss of function of the key homologous recombination genes BRCA1 and BRCA2 is associated with a range of distinct mutational signatures that include: 1) A single nucleotide variation based mutational signature (“COSMIC signature 3” or “BRCA signature” as labeled in the original publication[6]), 2) a short insertions/deletions based mutational profile, often dominated by deletions with microhomology, a sign of alternative repair mechanisms joining double-strand breaks in the absence of homologous recombination[7, 8] 3) large scale rearrangements such as non-clustered tandem duplications of a given size range (mainly associated with BRCA1 loss of function) or deletions in the range of 1-10kb (mainly associated with BRCA2 loss of function)[9]. We have recently shown that several of these DNA aberration profiles are in fact directly induced by the loss of BRCA1 or BRCA2 function[8].

Therefore, we analyzed all available whole genome sequencing data from the TCGA lung adenocarcinoma (LUAD) and squamous lung cancer (LUSC) cohorts and determined which of the above listed mutational signatures are present in these cases. Based on analyzing whole genome (n=42 and n=48, respectively) and whole exome (n=553 and n=489 samples) data we compared their utility for estimating HRD.

## Methods

The normal and tumor BAM files were downloaded for the whole exome sequencing (WXS) samples from TCGA. There were n=489 lung squamous carcinoma (LUSC) and n=553 lung adenocarcinoma (LUAD) samples available with both normal and tumor samples. The Mutect2 vcf files and the clinical data were downloaded from the TCGA data portal (portal.gdc.cancer.gov).

The BAM files for the whole genome sequenced samples were downloaded via the ICGC data portal (dcc.icgc.org). (LUSC: n=48, LUAD: n= 42 patients.)

### Mutation, Copy number, and Structural Variant Calling

Germline single nucleotide mutations were specifically called at and around the key HR-related genes for genotyping purposes using HaplotypeCaller, while somatic point-mutations and indels had been called using Mutect2 (GATK 3.8). In order to ensure the high fidelity of the reported SNVs, additional hard-filters had been applied to the resulting variants. In the germline case, the minimum mapping quality (PHRED) was set to 50, variant quality to 20 and a minimum coverage of 15 was ensured, while in case of the somatic SNVs and indels, the minimum tumor LOD (logarithm of odds) was set to 6, the normal to 4, the normal depth to 15, the tumor depth to 20 and the minimally allowed tumor allele frequency to 0.05.

Copy number profiles were determined using Sequenza[10], with fitted models in the ploidy range of [1,5] and cellularity range [0,1]. When a fitted model’s predictions significantly differed from the expected ploidy-cellularity values, an alternative solution was selected manually.

Structural variants were detected via BRASS (v6.0.0 - https://github.com/cancerit/BRASS). Through additional hard filters, the minimum amount of variant-supporting read-pairs was set to 6, and a successful local de novo assembly of the reads by velvet was demanded.

### Genotyping

The genotypes of the key homologous recombination related genes were determined via annotating the small-scale variant files using intervar[11]. Variants predicted as pathogenic or likely pathogenic were considered deleterious, while variants with unknown significance were treated with greater care but kept as wild-type.

### Mutational Signatures

Somatic point-mutational signatures were determined with the deconstructSigs R package[12], by using the cosmic signatures as a mutational-process matrix (Supplementary Figures 5-8).

The extraction of rearrangement signatures was executed according to the following strategy: first, the reported structural variants were mapped to the alphabet of the 32-dimensional structural variant-affecting mutational alphabet and stored into the matrices **M**_**LUSC**_ and **M**_**LUAD**_. Due to the low number of samples in the two WGS cohorts, the extraction of de novo rearrangement signatures was not achievable. Instead, a breast cancer-based, previously described matrix of mutational signatures (**P**) was used[9]. From these matrices, the signature composition (**E**) was estimated by solving the non-negative least squares problem||**P**\***E** – **M**||_2_, subject to E_ij_ > 0, for all i and j (Supplementary Figure 10).

### Genomic scar scores

The calculation of the genomics scar scores (loss-of-heterozygosity: LOH, large scale transitions: LST and number of telomeric allelic imbalances: ntAI) were determined using the scarHRD R package [13].

### HRDetect

Due to the absence of a sufficient number of authentic HR-deficient cases, the derivation of two separate, LUAD and LUSC specific HRDetect models was not achievable. Instead, on the whole genomes, the scores were calculated using the original, breast cancer-derived model, while the whole exomes relied on an alternative, whole exome-based, but also breast cancer-specific model (further details are available in the Supplementary Notes). In order to get to the results of Figure 2A, the whole genome features were log-transformed, and standardized within their respective cohorts (i.e. N_LUSC = 49, N_LUAD = 42), i.e. the z-scores of each of the log-transformed attributes had been calculated using the means and standard deviations of the 49 LUSC and 42 LUAD samples respectively. Since the original model was trained on 560 breast cancer whole genomes, the standardization step was executed using the means and standard deviations of the 560 breast cancer whole genome as well. The distributions of these alternative (referred to as “breast cancer standardized”) HRDetect predictions are available in Supplementary Figure 11, and both the scores and HRDetect predictor attributes are available in Supplementary Tables 1 and 2.

#### HRDetect values in cell lines

The mutational information for lung adenocarcinoma and squamous cell carcinoma cell lines were obtained from the CCLE [14], PARP-inhibitor sensitivity was downloaded from the GDSC project [15]. The mutational signatures were determined using deconstructSigs R package. Due to the missing reliable allele-specific copy number data, a simplified HRDetect was applied using Signature 3,6,8,13,17, and proportion of microhomology-mediated deletions (Further details: Supplementary Notes.)

## Results

### Loss of function mutations of HR genes in lung cancer

Detailed analysis on the germline and somatic mutations of DNA repair genes were performed. We identified loss of function mutations for the BRCA2 gene in three LUSC and two LUAD cases (all three in LUSC and one in LUAD were coupled with LOH, however, in the tumor of the LUSC donor: TCGA-66-2744, the allele harboring the germline BRCA2 mutation was lost due to an LOH), and loss of function mutation for BRCA1 for one case in LUSC (Supplementary Figures 1-4). A LUAD case was also identified with a RAD51B germline mutation that was recently shown to be associated with HR deficiency[16] accompanied with an LOH in the tumor. We hypothesized that some of these cases may exhibit robust signs of homologous recombination deficiency induced mutational signatures.

### HR deficiency associated mutational signatures in lung squamous carcinoma

The two BRCA2 mutant LUSC samples exhibited an elevated short deletion/insertion ratio of >2 with one of them having the highest such ratio in the cohort (Figure 1A). Increased deletion/insertion ratios were described previously for BRCA2 deficient cancers using whole genome sequencing data[17]. The same cases also showed the highest proportion (>0.1) of larger than 2 bp long microhomology mediated deletions and the highest proportion of larger than 9 bp long short deletions (Supplementary Figure 9). These two indel-patterns have also been described previously in BRCA2 deficient human cancer biopsies[7, 8, 17].

**Figure 1.**
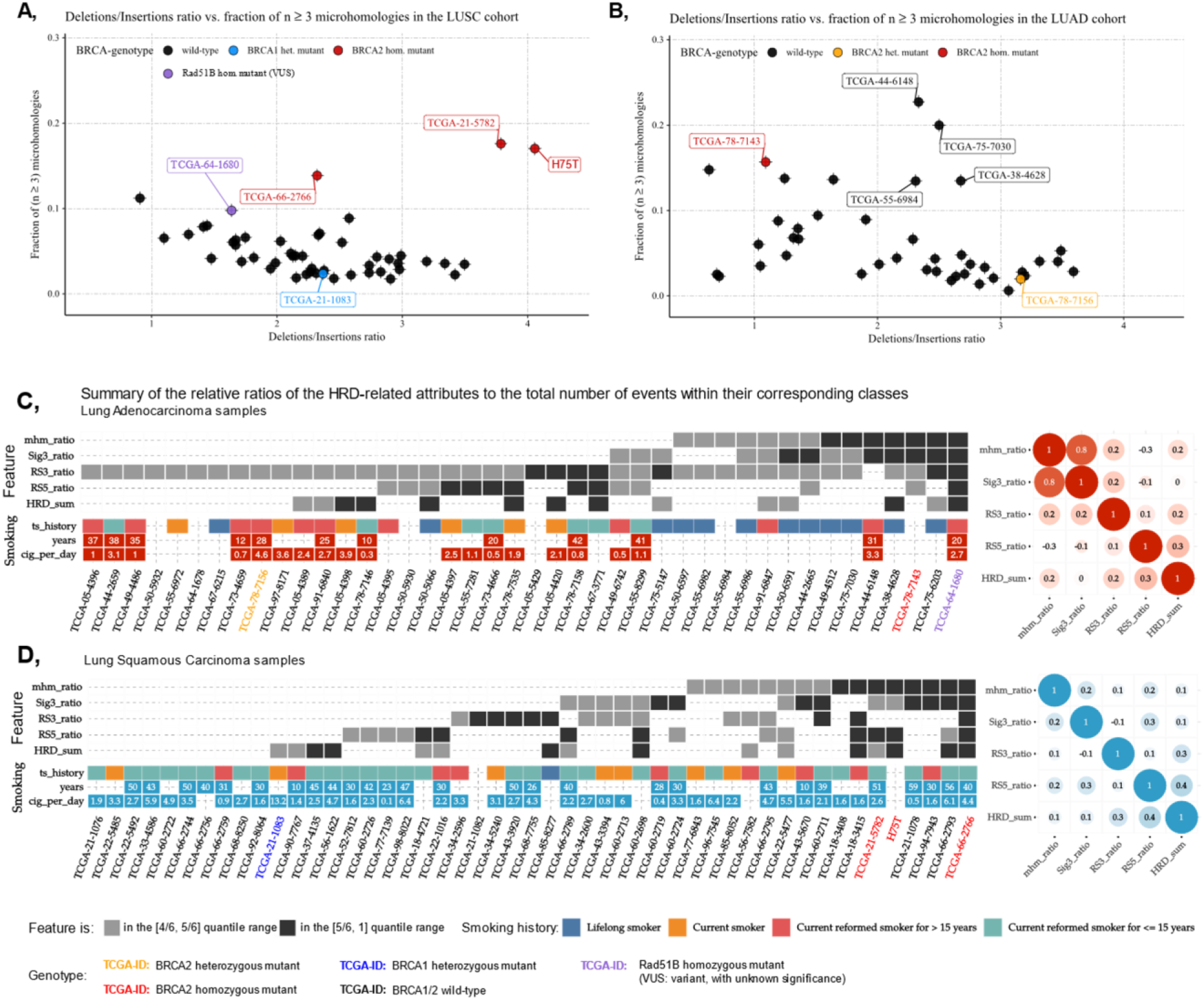
Summary of the HRD-related predictors in the LUAD and LUSC whole genome dataset. **A)** Fraction of microhomology-mediated deletions with larger or equal to 3 bp in length, versus the deletions/insertions ratio in the TCGA-LUSC cohort. **B)** Fraction of microhomology-mediated deletions with larger or equal to 3 bp in length, versus the deletions/insertions ratio in the TCGA-LUAD cohort. **C,D)** Summary of the HRD-related molecular features extracted from the TCGA-LUAD (C) and TCGA-LUSC (D) whole genome samples. The considered features are as follows: ***HRD_sum***-the sum of the three allele-specific CNV-derived genomic scars scores (HRD-LOH + LST + ntAI), **RS3_ratio**: Rearrangement signature 3 ratio, i.e. the number of structural variants originating from Rearrangement Signature 3, divided by the total number of structural variants in the sample, **RS5_ratio:** Rearrangement Signature 5 ratio, i.e. the number of structural variants originating from rearrangement signatures 5, divided by the total number of structural variants in the sample, **mhm_ratio:** the ratio of microhomology-mediated deletions, i.e. the number of deletions, that begin with at least three consecutive nucleotides that it shares with the sequence that follows the deletion, divided by the total number of deletions, **Sig3_ratio:** Substitution signature 3 ratio, i.e. the number of somatic point mutations that can be contributed to the activity of the molecular process captured by Substitution Signature 3, divided by the total number of single nucleotide somatic mutations. The top left sections of the two panels contain dark and light gray tiles, which represent the positions of the samples within the distributions of each of the above-described features. A dark grey tile means, that the corresponding ratio (vertical axis) of the sample (horizontal axis) is among the highest within the cohort; it lies between the 5^th^ and 6^th^ sextiles, i.e. above the 83rd percentile. A light grey tile indicates that the corresponding feature-ratio is between the 4^th^ and 5^th^ sextiles, i.e. between the 66^th^ and 83rd percentiles. The bottom section of the panels shows the available tobacco history of the samples. Empty tiles either indicate that at the time of data collection the donor was registered as a non-smoker, or that the information was not available. To the right sides of the two panels, two correlograms show the Pearson correlation coefficients between all the ratio pairs. Apart from the correlation coefficient between the Signature 3 and microhomology-mediated deletion ratios within the LUAD cohort, all other coefficients were mediocre or negligibly small.

We did not detect an increased SNV signature 3 (originally described in BRCA1/2 deficient tumors[6]) in these particular cases, probably because the high level of smoking induced mutational signatures would mask them even if they were present (Figure 1C and D). (For a detailed distribution of SNV based signatures see Suppl. Fig 8).

On the other hand, both of the above-mentioned BRCA2 deficient cases showed the highest number of RS5 rearrangement signatures (Figure 1D), which were previously described in breast cancer to be strongly associated with loss of function of BRCA2[9].

Taken together, both of the likely BRCA2 deficient LUSC specimens showed clear signs of BRCA2 deficiency associated mutational signatures and thus those cases are likely homologous recombination deficient.

### HR deficiency associated mutational signatures in lung adenocarcinoma

In the case of LUAD, consistent with the lower number of smokers in this tumor type, in about half of the cases (20 out of 42) the smoking signature did not dominate the SNV signatures and the contribution of other mutational processes could be clearly detected (Supplementary Figure 8). More prominently, the BRCA2 mutant case with LOH (TCGA-78-7143), along with six other cases, showed a strong presence of signature 3. The same BRCA2 mutant sample had a high proportion of microhomology-mediated deletions and four of the other six samples showed high deletion/insertion ratios along with high proportions of microhomology-mediated deletions (Figure 1B). The RAD51B case (TCGA-64-1680) showed both the signs of the HR deficiency associated indel patterns and a high signature 3 ratio.

The BRCA2 mutant case and the four other cases showing HRD-like SNV and indel patterns also showed presence of the rearrangement signatures associated with BRCA function loss, although to a lesser extent than that seen in TCGA-64-1680 and in BRCA1/2 mutant breast cancer in general.

### HRDetect scores in the LUAD and LUSC WGS cohorts

Considering the different types of mutational signatures induced by the loss of function of HR genes, and that in a given tumor the loss of that gene may have a different impact on those mutational signatures, it was suggested recently that the SNV, short indel and large-scale rearrangement signatures along with a CNV-derived genomic scar score[18] be combined into a single HRD quantifier, HRDetect[19]. This complex HR deficiency measure was trained on the number and relative distribution of HRD induced DNA aberration profiles in breast cancer.

We calculated the breast cancer trained HRDetect values for all WGS cases by standardizing the lung predictors combined with the original breast cancer dataset, and found that the two above described, likely BRCA2 deficient LUSC cases; TCGA-66-2766 and TCGA-21-5782 have the highest HRDetect values, the former of which even exceeded 0.7, which was proposed to be the threshold value for bona fide HR deficient cases in breast cancer (Supplementary Figure 11).

We also calculated the HRDetect values when the predictors were standardized on the lung cancer cases alone (Figure 2A). These values are in general higher since, as we pointed out, some of the HRD suspect cases showed strong signs of some (e.g. SNV and short indel based) HRD signatures but not others (large rearrangement based signatures) (Figure 1C and D). In other words, it is possible that the individual parameters in a lung cancer specific HRDetect model will be significantly different from those in breast cancer. Both in the case of LUAD and LUSC, eight of the analyzed cases showed a >0.7 lung cancer normalized HRDetect value (Figure 2A) estimating the number of HR deficient cases at less than 20%.

**Figure 2.**
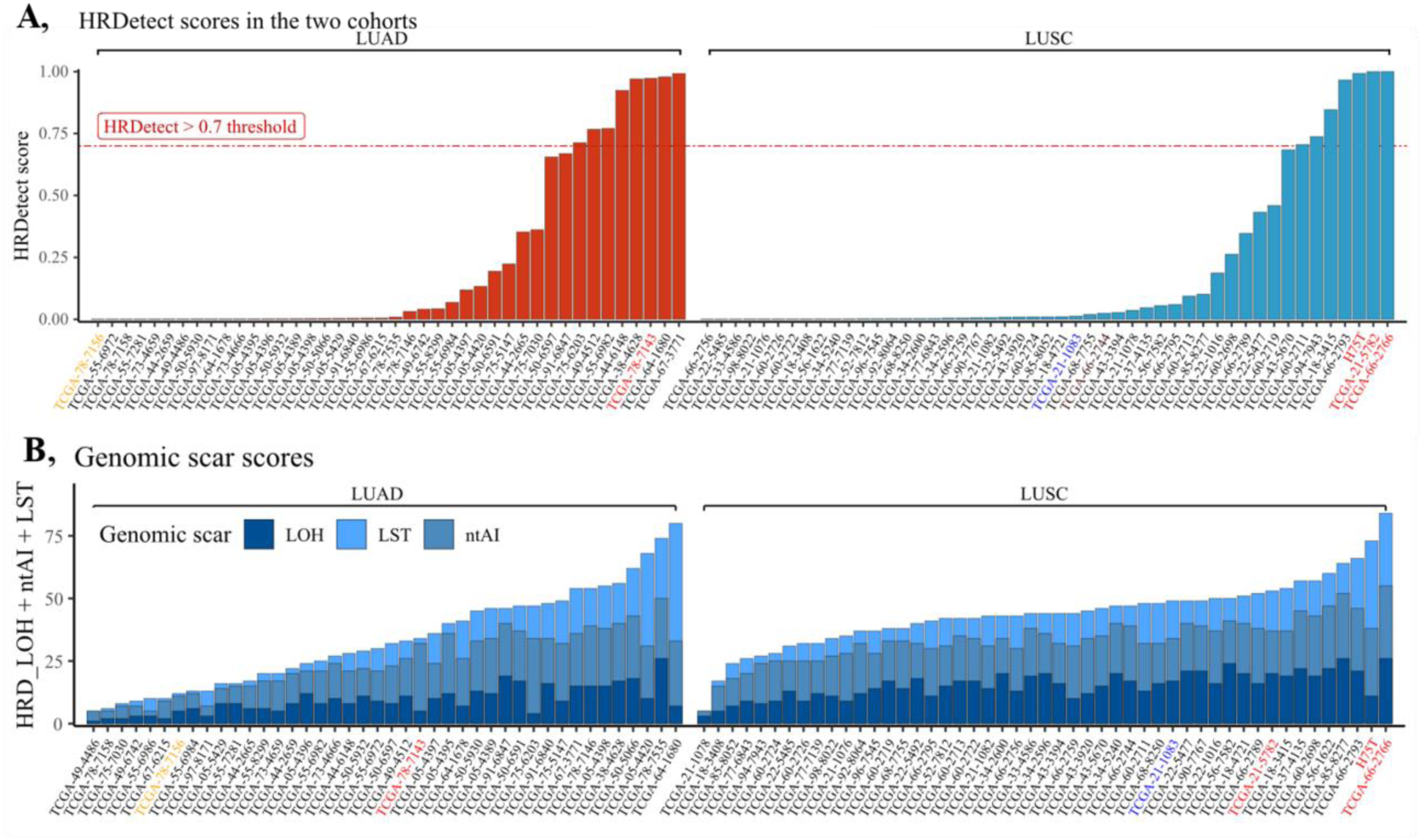
HRD scar scores and HRDetect scores of the LUAD and LUSC WGS datasets. In both panels, the sample names are colored according to their genotypes: yellow – BRCA2 heterozygote mutant, red – BRCA2 homozygote mutant, blue – BRCA1 heterozygote mutant. **A)** HRDetect scores calculated using the original, breast cancer whole genome-based HRDetect weights by following the original article’s standardization and attribute-transformation strategies [19]. **B)** The total sum of the genomic scar scores (HRD-LOH, LST, and ntAI) determined from the LUAD and LUSC whole cancer genome’s allele-specific copy number profiles.

### HR deficiency associated biomarkers in whole exome sequencing data

While whole genome sequencing data carry the most information about HRD induced mutational processes, we previously showed that in the case of breast cancer whole exome sequencing data can also be used for these purposes, albeit using only 1 % of all aberrations that are present in the whole genome data[20]. We wanted to determine how well WES data capture HR-deficiency induced mutational signatures in lung cancer.

We started with the comparative analysis of those cases that had both WGS and WES data available. (Suppl Figure 14). For the LUSC cases, the HRD-LOH score showed strong (0.83), and the ratio of signature 3 showed reasonable (0.41) correlation across the WES and WGS data. Due to the lower number of detectable deletions in whole exomes in general, all microhomology-mediated deletions were considered in WES if their size exceeded 1bp, and this number was compared to the >2bp microhomologies in whole genomes. While on average there was a two order of magnitudes difference in the absolute number of deletions between the corresponding pairs, they exhibited a strong correlation (0.79). Of the two most likely HR deficient LUSC cases, TCGA-21-5782 showed good correlation of all three measures across the WES and WGS data. In the other case (TCGA-66-2766), however, the WES data did not recapture the same mutational signatures as the WGS data (Supplementary Figure 13).

For LUAD all three measures showed correlations between 0.55 and 0.84 across the WGS and WES data, and the cases with high HR deficiency associated attributes, like signature 3 or the HR deficiency-like insertion/deletion pattern in the WGS data had shown the same tendency in WES data as well.

For the distribution of the various HR-deficiency mutational signatures across the entire WES based cohorts see Supplementary Figures 12 and 13.

### HRDetect scores in the LUAD and LUSC whole exome sequencing data

We calculated the HRDetect values based on the standardized and log-transformed attributes of the WES data (further details of the whole exome model are available in the Supplementary Notes). We found only moderate correlations (~0.5) between the corresponding pairs in the lung cancer cohorts (Supplementary Figure 14). This limited correlation may be the reason for the lack of enrichment of BRCA1/2 deficient cases in the WES cohorts (Supplementary Figure 15).

We estimated the number of high HRDetect values in the larger, WES cohorts as well and those numbers were lower than those detected in the WGS data. (It was about 16% for LUSC and 4% for LUAD)

Since high HRDetect scores were reported to be associated with better clinical outcome in platinum treated breast cancer[21], we were wondering whether lung cancer cases below and above the HRDetect thresholds that we determined in the WES data have significantly different outcome when treated with platinum-containing therapy. However, higher HRDetect-scores were not associated with better outcome in these cohorts. (Supplementary Figures 16 and 17).

### HR deficiency associated mutational signatures in a lung squamous cell carcinoma case with exceptionally good response to platinum treatment

In order to validate the clinical utility of HRD associated mutational signatures in lung cancer we searched our clinical database for advanced lung cancer cases with exceptionally good response to platinum-based therapy and available fresh frozen material for whole genome sequencing. We identified a stage IVA lung squamous carcinoma case (see case description in the Supplementary Notes) with a durable (> 20 months), symptom-free survival in response to platinum-based treatment (H75T). Since the patient was a heavy smoker the SNV based mutational signatures were dominated by the smoking-associated SNV signature 4 (Supplementary Figure 8). On the other hand, this tumor showed all short indel consequences of HRD: A high deletion/insertion ratio, high fraction of microhomology-mediated deletions and high fraction of larger than 10 bp deletions (Figure 1 and Supplementary Figure 9). It also had a high number of BRCA2 deletion associated large scale rearrangement events (1-10 kb deletions)(Supplementary Figure 10). Finally, the combined HRD measures, the HRD-LOH and HRDetect scores, ranked amongst the highest we analyzed from TCGA (Figure 2B). Finally, the germline DNA analysis uncovered a loss of function BRCA2 mutation that, in combination with a somatic LOH, likely induced the HRD associated mutational profiles (Supplementary Figures 2-4).

### HRDetect scores of lung cancer cell lines are correlated with PARP inhibitor sensitivity

We calculated the HRDetect scores for 67 lung cancer cell lines for which both sequencing data and drug sensitivity data were available in the CCLE and GDSC databases. Most (n=58, 87%) cell lines had a low HRDetect value <0.25). Eight cell lines had a >0.70 HRDetet score and these were significantly more sensitive to olaparib and talazoparib treatment (Supplemenary Figure 18).

## Discussion

PARP inhibitors show significant clinical efficacy in tumor types that are often associated with BRCA1/2 mutations, such as breast, ovarian, and prostate cancer. In order to further expand this clinical benefit, there are several ongoing clinical trials evaluating the efficacy of PARP inhibitors in non-small cell lung cancer, such as the PIPSeN (NCT02679963) and Jasper (NCT03308942) trials. If, however, the clinical benefit is strongly associated with HRD in this tumor type and only a minority of lung cancer cases harbor this DNA repair pathway aberration, then the success of those clinical trials will greatly depend on our ability to identify and prioritize the HRD cases.

In order to develop such a diagnostic method, we first analyzed the BRCA1/2 mutant lung cancer cases. Lung cancer is usually not associated with germline BRCA1/2 mutations, although sporadic cases have been reported[5, 22].Nevertheless, due to e.g. smoking, about 5-10% of non-small cell lung cancer cases show somatic mutations in either the BRCA1 or BRCA2 genes. Some of those are likely to be pathogenic and associated with LOH as well [5]. In our analysis, these cases, when analyzed by whole genome sequencing, clearly showed the mutational signatures usually associated with HRD. This strongly suggests that there are some bona fide HRD cases amongst lung cancer as well. Beyond mutations in BRCA1/2, HRD can be induced by a variety of mechanisms, such as suppression of expression of BRCA1 by promoter methylation. This is reflected by the fact that a significant number of ovarian and breast cancer cases show clear patterns of HRD associated mutational signatures in the absence of mutations of BRCA1/2 or other key HR genes[19]. Furthermore, BRCA1 mutant cases can be rendered HR proficient and thus PARP inhibitor resistant by the loss of other genes such as 53BP1 or REV7, etc[23, 24]. Therefore, downstream mutational signatures, such as those investigated in our analysis, could be more accurate measures of HRD than the mutational status or expression change of individual genes and they could serve as a complementary biomarker to the mutation status of HR associated genes In fact, we identified several lung cancer WGS cases with high HRD induced mutational signatures that were not associated with BRCA1/2 mutations and those could also be responding to PARP inhibitor therapy as our cell line based preclinical analysis suggests.. We made every effort to detect a likely explanation for the cases with significant HRD associated mutational signatures but TCGA profiles have significant limitations due to e.g. normal tissue contamination. For example, significant expression deficiency or LOH of the BRCA1 or BRCA2 genes can often be masked by the presence of these genes in the normal cells in the tumor biopsy.

It is important to estimate the proportion of potentially HRD non-small lung cancer cases to optimize PARP inhibitor trials. Using the limited number (less than one hundred in total), WGS covered lung cancer cases we estimated the proportion of truly HRD cases as less than 20%. In our analysis, WES data seemed to be less accurate to determine HRD status in lung cancer. This could be due to, e.g. the high level of smoking induced mutational signatures interfering with the HRD induced mutational signatures. The limitations of WES based HRD classifiers were also highlighted in a recent publication that WES profiled several BRCA1/2 deficient NSCLC cases [5]. In these cases, in accordance with our results, the SNV based mutational signatures were dominated by the smoking signature and the composite HRD score of these BRCA1/2 deficient NSCLC cases was not higher than that of the control cases. As we pointed out, HRD associated indels and large-scale rearrangement signatures are key components of HRD measures but those can be detected with much lower accuracy from WES than from WGS data. When we investigated the mutational signatures from whole genome sequencing data of an exceptionally good responder of lung cancer to platinum-based therapy, the high levels of HRD associated mutational signatures were apparent, despite the fact that the patient was a smoker. This strongly suggests that bona fide HR deficient cases with clinical consequence can be identified by WGS analysis but not necessarily by WES analysis, warranting further, WGS based clinical investigations.

We used the WES derived HRD measures to investigate the correlation between the likely presence HRD and better survival upon treatment in lung cancer in TCGA. We did not see any correlation which is probably due to a number of factors. These patients were treated in addition to platinum with other agents as well. Sensitivity or resistance to platinum treatment is also associated with several other mechanisms in addition to HRD [25, 26] and the WES derived HRD associated mutational signatures are less accurate than their WGS based counterparts.

## Acknowledgment

This work was supported by the Research and Technology Innovation Fund (KTIA_NAP_13-2014-0021 and NAP2-2017-1.2.1-NKP-0002 to Z.S.); Breast Cancer Research Foundation (BCRF-17-156 to Z.S.) and the Novo Nordisk Foundation Interdisciplinary Synergy Programme Grant (NNF15OC0016584 to Z.S. and I.C.), The Danish Cancer Society grant (R90-A6213 to MK). The results shown here are based upon data generated by the TCGA Research Network: http://cancergenome.nih.gov/.

## Conflict of Interest

Z. Szallasi is an inventor on a patent used in the myChoice HRD assay.

## Authors contribution

MD, ZS and JB designed the analysis and performed all computational analysis and participated in preparation of the manuscript. MK, VT, SS, OR, IC, JM, AGP, and DS participated in designing the analysis and prepared the manuscript. ZS designed the analysis, had supervisory role and prepared the manuscript.

