## Supplementary Notes for "A subset of lung cancer cases shows robust signs of homologous recombination deficiency associated genomic mutational signatures"

Supplementary Material

### Contents

### List of Figures

### Supplementary tables not included in the Supplementary Text

Supplementary Table 1-2 are available separately, in csv format.

- **Supplementary Table 1**: List of the LUAD whole genomes with their HRD-related attributes
- **Supplementary Table 2**: List of the LUSC whole genomes with their HRD-related attributes

### 1 Analyzed cohorts

#### 1.1 Whole Genomes

42 LUAD and 48 LUSC WGScohorts had been downloaded from the icgc data portal:

- LUAD cohort: https://icgc.org/ZV9
- LUSC cohort: https://icgc.org/ZVC

#### 1.2 Whole Exomes

Both binary alignment files and MuTect2 vcfs had been downloaded from the gdc data portal. Altogether 553 LUAD and 489 LUSC whole exomes were considered.

#### 1.3 Donors with both WGS and WES samples

The majority of the patients who had whole genome data available, had whole exome sequences as well. All the 48 patients in the LUSC WGS cohort had at least 1 corresponding whole exome, however out of the 42 LUAD WGS patients only 39 had whole exomes. The WGS samples without pairs:

- TCGA-05-5429
- TCGA-64-1678
- **TCGA-78-7143**

Since **TCGA-78-7143** had a likely pathogenic germline BRCA2 mutation coupled with an LOH, we have created an exonic bam-slice using the reads that cover the exome from the WGS bam, in order to check whether the BRCAness phenotype is detectable in the exonic version.

### 2 Genotyping

Genotypes were determined according to the following scheme; we have called germline variants via GATK (v3.8) HaplotypeCaller, and on whole genomes somatic variants with GATK (v3.8) MuTect2 (for whole exomes, MuTect2-derived vcfs were already available from the gdc data portal). The pathogenecity of these variants was assessed using Intervar (v2.0). From the resulting variant files only the exonic and the +/- 10 nucleotide regions around the exons (in order to account for the possible splice-variants) were considered. From these mutations only those were kept, that were predicted as "Likely Pathogenic", "Pathogenic" or "Uncertain" according to ClinVar (20170905). At last, a threshold on the depth of these variants were set to 20.

If a variant had been characterized as pathogenic or likely pathogenic by intervar, the corresponding sample was considered mutant, assuming that at least a heterozygous mutation is present in the sample. Variants with unknown significance were collected separately, but they did not affect the genotyping scheme.

### 3 Case report of the platinum sensitive LUSC patient: H75T

A 76-year-old man was diagnosed with solitary pulmonary nodule in the right upper lobe during a screening chest X-ray in August, 2016. He was an ex-smoker with 35 pack-year index, and had quitted smoking 20 years ago. He suffered from atherosclerosis and cardiovascular disease. In January, 2017 segmental surgical resection of the right upper lobe was performed with a pathological diagnosis of poorly differentiated squamous cell lung carcinoma of 31 mm in diameter with lymphoid vessel invasion (pT2a-N0-M0). He received no adjuvant oncotherapy. In June, 2017 PET-CT revealed bilateral pulmonary dissemination (of maximum 8x16 mm) and enlargement of the mediastinal lymph nodes with (18)F-FDG avidity, therefore, gemcitabine-carboplatin chemotherapy was indicated by a multidisciplinary tumor board. After 4 cycles of this treatment partial response could be observed using the RECICST 1.1 criteria. In March, 2018 no evidence of tumor could be demonstrated on chest CT. In April, 2019 the patient is still alive.

##### 3.1 Lung adenocarcinoma WGS samples - mutations in HR-relevant genes

Mutations found in the LUAD WGS cohort are summarized in Suppl.Fig. **??**.

##### 3.2 Lung squamous carcinoma WGS samples - mutations in HR-relevant genes

Mutations found in the LUSC WGS cohort are summarized in Suppl.Fig. **??**.

##### 3.3 Loss of Heterozygosity

The occurrance of Loss of heterozygosity was estimated using the samples' sequenza-derived copy-number segments. If the copy-numbers of either the A or B alleles dropped to zero within the coordinates of a gene, then the LOH event was registered (Suppl. Fig. **??**).

##### 3.4 Methylation

Since the majority of the samples had only HumanMethylation 27k data available or didn't have methylation info at all, the genotyping scheme did not consider the methylation status of the gene-specific probes.

##### 3.5 Final genotypes

The final genotypes are summarized in Suppl.Fig. **??**. Both cohorts had likely pathogenic heterozygous or homozygous BRCA1/2 mutants among their samples:

LUAD:

- TCGA-75-7156 (likely pathogenic BRCA2 germline mutation)
  /frameshift insertion at chr13:32912949,T>TTGTGC/
- TCGA-78-7143 (likely pathogenic BRCA2 germline mutation + LOH)
  /frameshift insertion at chr13:32906473, A>ACCTAATCTTACTATAT/

LUSC:

- TCGA-21-1083 (likely pathogenic BRCA1 somatic mutation)
  /stopgain SNV at chr17:41244585, G>C/
- TCGA-21-5782 (likely pathogenic BRCA2 somatic mutation + LOH)
  /frameshift deletion at chr13:32930627, AG>A/
- TCGA-66-2744 (likely pathogenic BRCA2 germline mutation + LOH)
  /frameshift deletion at chr13:32912337, CTG>C/
- TCGA-66-2766 (likely pathogenic BRCA2 germline mutation + LOH)
  /stopgain SNV at chr13:32914349, G>T/

In addition, TCGA-64-1680 – a LUAD sample with high HRD-related genomic aberration scores – had a UNK germline mutation in RAD51B /nonsynonymous SNV at chr14:68352672, A>G/.

##### 3.6 Whole Exomes

Genotyping of the whole exomes followed a similar strategy to the whole genomes'.

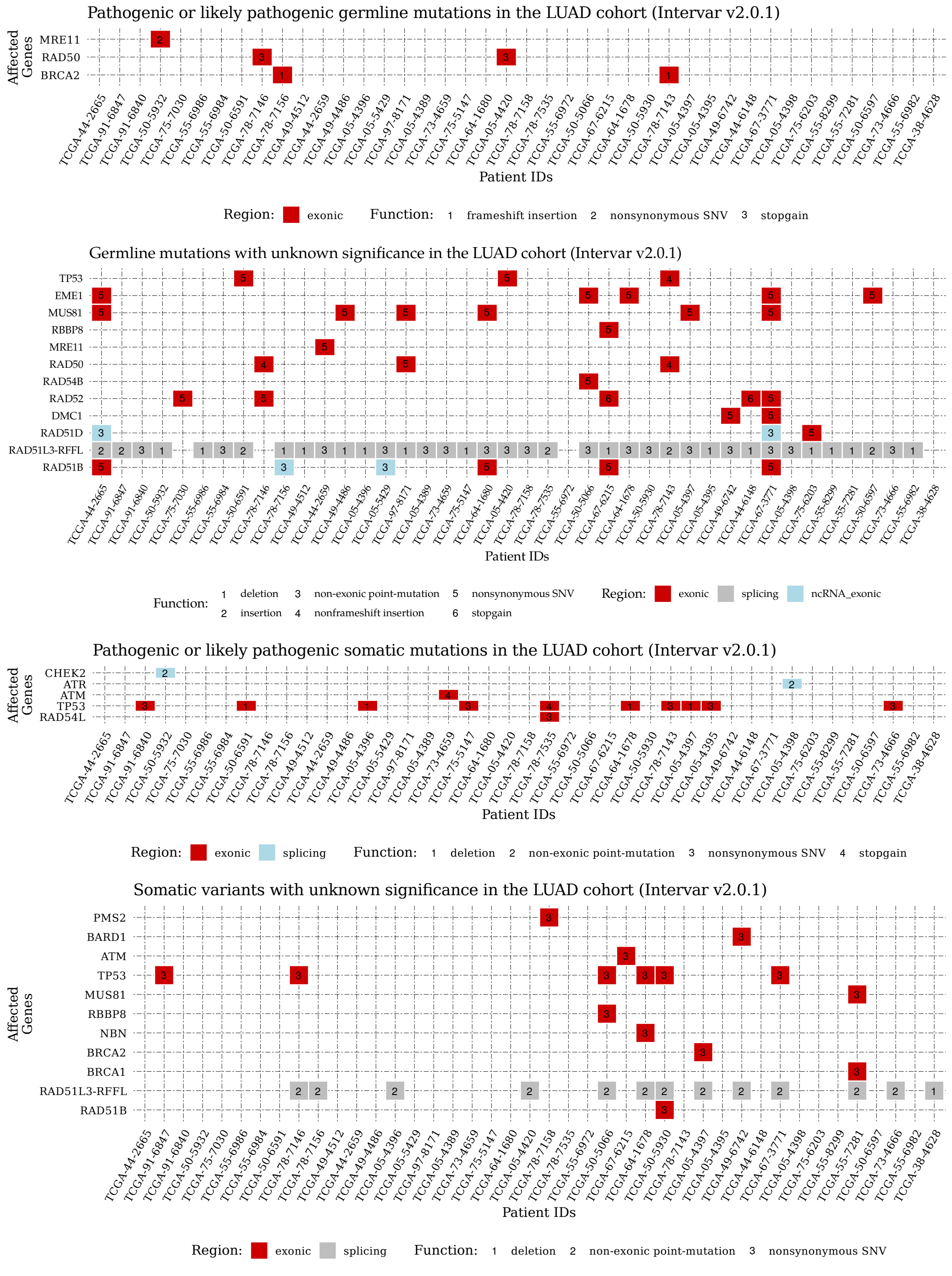


**Suppl.Fig. 1:** *First two panel from the top: Pathogenic or likely pathogenic and UNK germline mutations in the LUAD WGS cohort. Bottom two panel: Somatic variants in the LUAD WGS cohort*

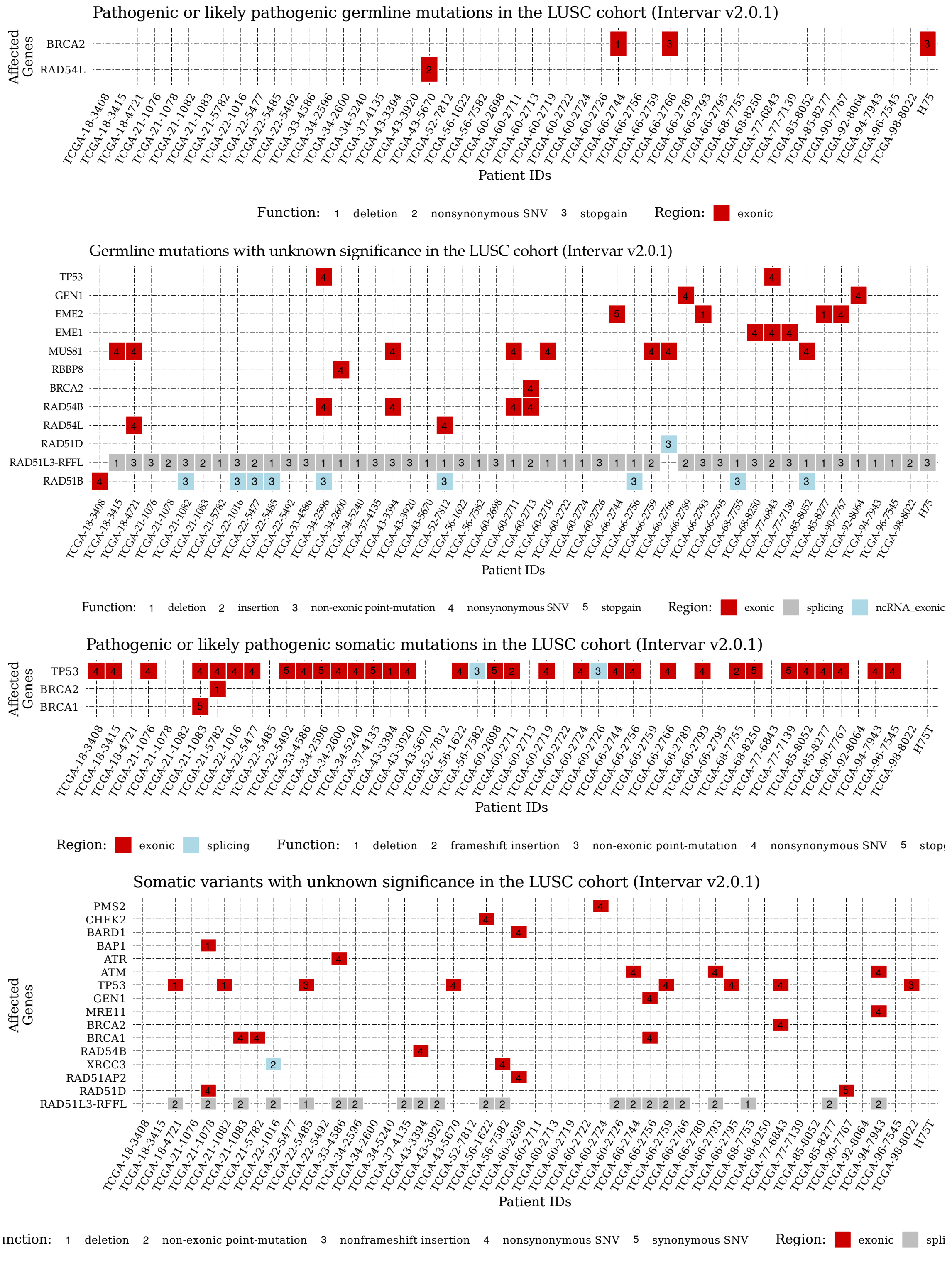


**Suppl.Fig. 2:** *First two panel from the top: Pathogenic or likely pathogenic and UNK germline mutations in the LUSC WGS cohort. Bottom two panel: Somatic variants in the LUSC WGS cohort*

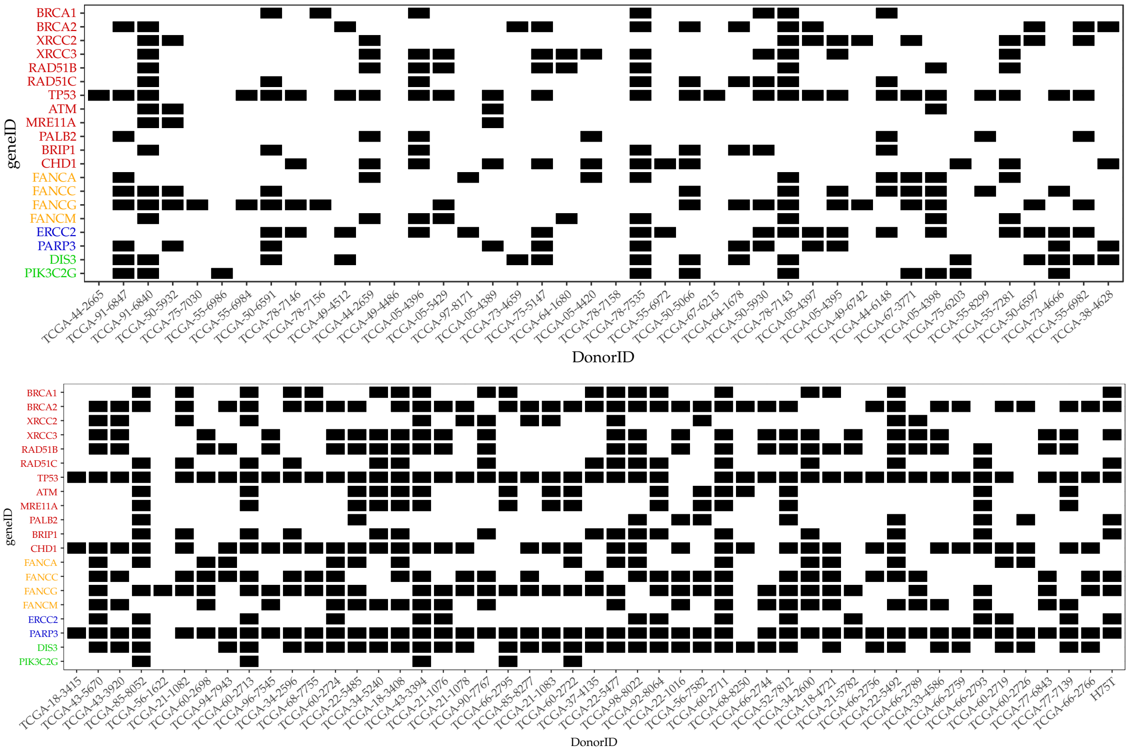


**Suppl.Fig. 3:** *Estimated occurrences of LOH events in the analyzed genes. Segment means are estimated using the sequenza and copynumber R packages.*

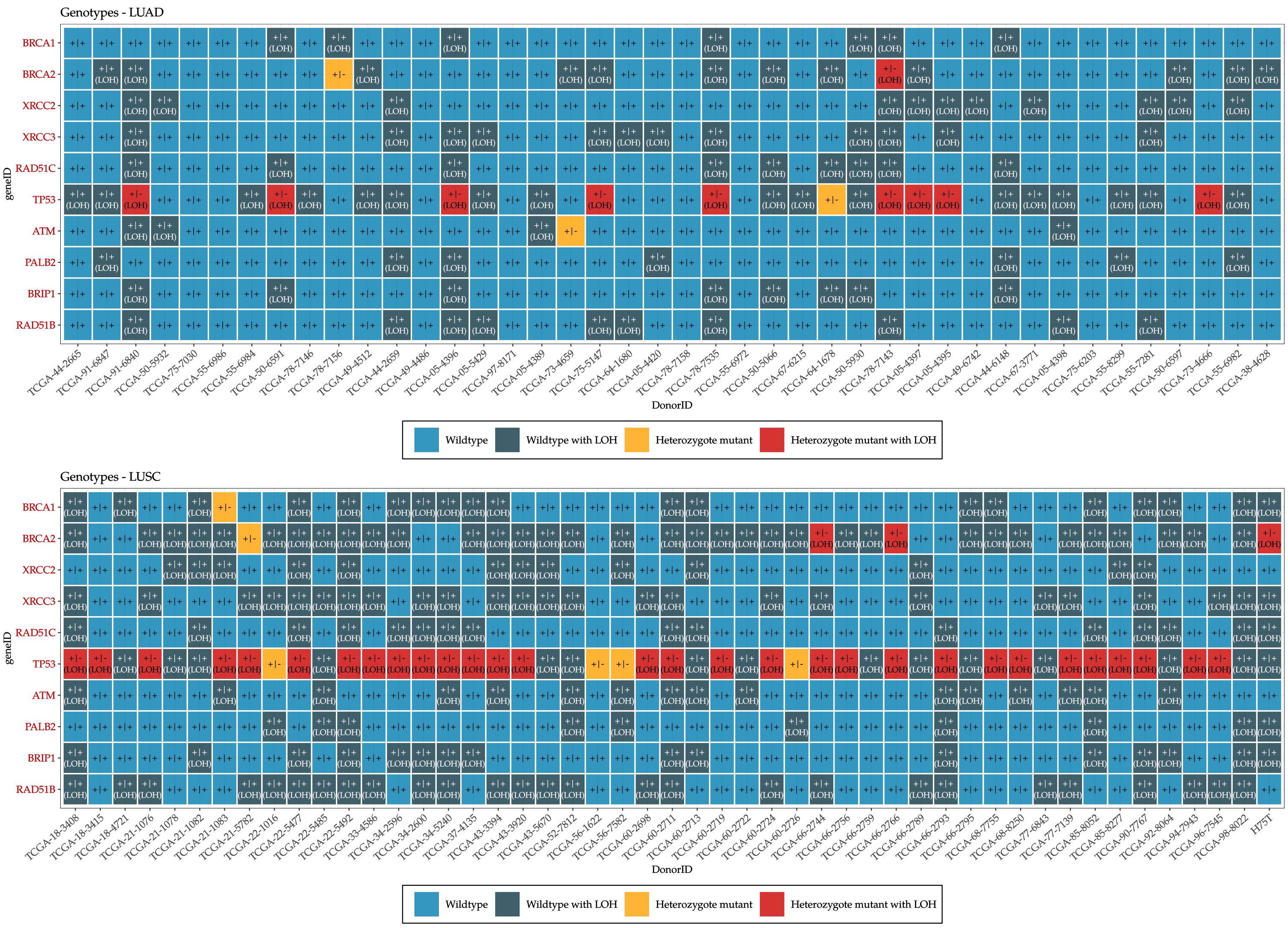


**Suppl.Fig. 4:** *Final genotypes of the LUAD and LUSC WGS cohorts. Genotyping is based on the presence of a pathogenic or likely pathogenic somatic/germline mutation in the gene and whether a loss of heterozygosity event accompanies them. - Heterozygote: at least a germline/somatic mutation present, but no LOH, homozygote: at least a germline/somatic mutation present AND an LOH.*

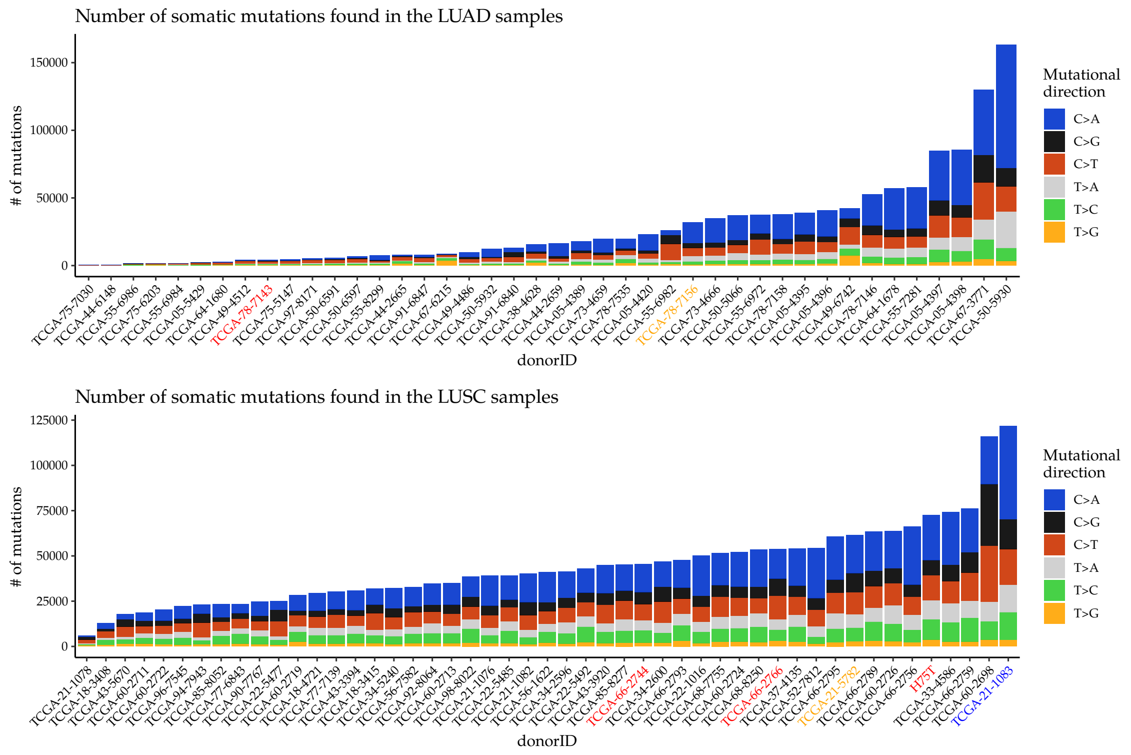


**Suppl.Fig. 5:** *Summary of the somatic substitutions detected in the whole genome cohorts. The vertical axis contains overall numbers, the colors on the bars indicate the relative composition of the mutational directions. Most samples are dominated by C>A mutations. Top panel: LUAD, Bottom panel: LUSC.*

### 4 HRD-related biomarkers

The HRD-induced genomic fingerprints analyzed in this study were the following:

1. Somatic Substitution Signatures [**?**]
2. Microhomology-mediated deletion ratio, and insertion/deletion ratio [**?**, **?**]
3. Genomic scar scores [**?**, **?**, **?**]
4. Rearrangement Signatures [**?**]

#### 4.1 Somatic Substitution Signatures

Somatic variants returned by MuTect2 had to PASS the following criteria as well, on top of the default filters of MuTect2

- TLOD $\geq 6$
- NLOD $\geq 3$
- Normal depth $\geq 15$
- Tumor depth $\geq 20$
- Alt. allele supporting reads in the tumor $\geq 5$

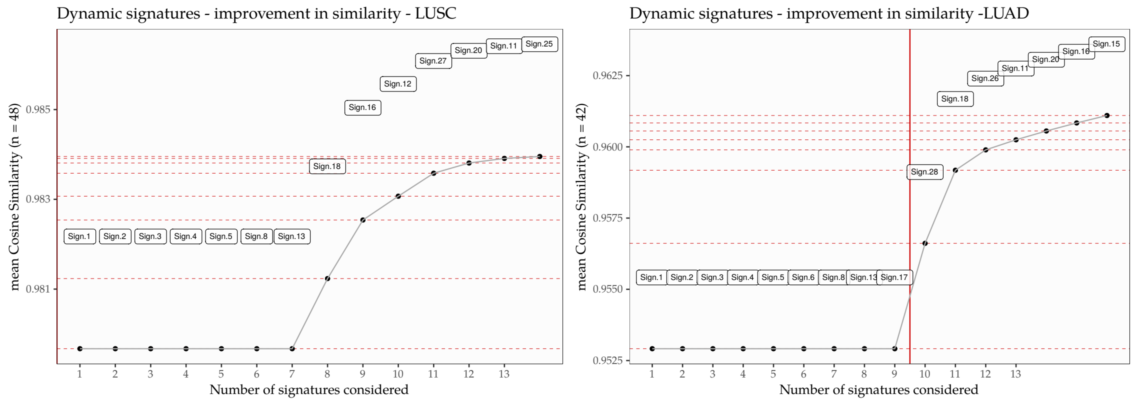


**Suppl.Fig. 6:** *A graphical representation of the dynamic signature addition process. In each iteration a signature that would improve the most on the cosine similarities were considered, however no additional signatures had been added to the list in either cohort. Left: LUSC, Right: LUAD*

- Alt. allele supporting reads in the normal = 0
- Alt. allele frequency $\geq$ 0.05
- FILTER field = "PASS"

The resulting distribution of SNV is displayed in Suppl. Fig. **??**.

Somatic signatures were extracted with the help of the `deconstructSigs` R package [**?**]. The list of possible mutational processes whose signatures' linear combination could lead to the final mutational catalogs (a.k.a. mutational spectra) was confined to those, that were reportedly present in lung adenocarcinomas and squamous carcinomas according to the COSMIC database (i.e. in LUAD: Signatures 1, 2, 4, 5, 6, 13, and 17, in LUSC: 1, 2, 4, 5, and 13). Furthermore, since we were primarily interested in their HR-related signature composition, we have added Signature 3 and 8 to the lists. After the evaluation of their signature compositions, the mutational catalogs of the samples were reconstructed, and the cosine of the angle between the 96-dimensional original and reconstructed vectors were measured (cosine similarity). Using this technique, we have also checked whether the incorporation of any additional signatures would improve the mean reconstruction similarities significantly, but the improvement was negligible in both WGS cohorts (Suppl. Fig. **??**). In general, the final cosine similarities were adequately high, especially between the original and reconstructed squamous carcinoma whole genomes (Suppl. Fig. **??**).
The final mutational signatures can be observed in Suppl. Fig. **??**.

#### 4.2 Classification of deletions

It has been shown recently, that cancer cells that exhibit homologous recombination deficiency, have unique characteristics in their indel profiles. Specimens with biallelic BRCA1/2 mutations have significantly more deletions that are longer than 10 bp than BRCA1/2 wild-type tumors, and they also tend to have more deletions than insertions [**?**]. It has also been found, that these deletions mostly arise due to the activity of the Microhomology Mediated End Joining (MMEJ) or the Single Strand Annealing (SSA) DNA repair pathways, and thus the relative ratio of microhomology mediated (mhm) deletions among them is significantly higher than in HR-competent cases [**?**]. Since the HR and MMEJ pathways differentiate at the point when RPA binds to the ssDNA overhangs, a dysfunctional BRCA2 protein involuntarily gives rise to an increased MMEJ/SSA activity. Non-surprisingly, the aforementioned increase in the mhm-deletion ratio is much more obvious in samples with BRCA2-/- mutations than in BRCA1-/- tumors.

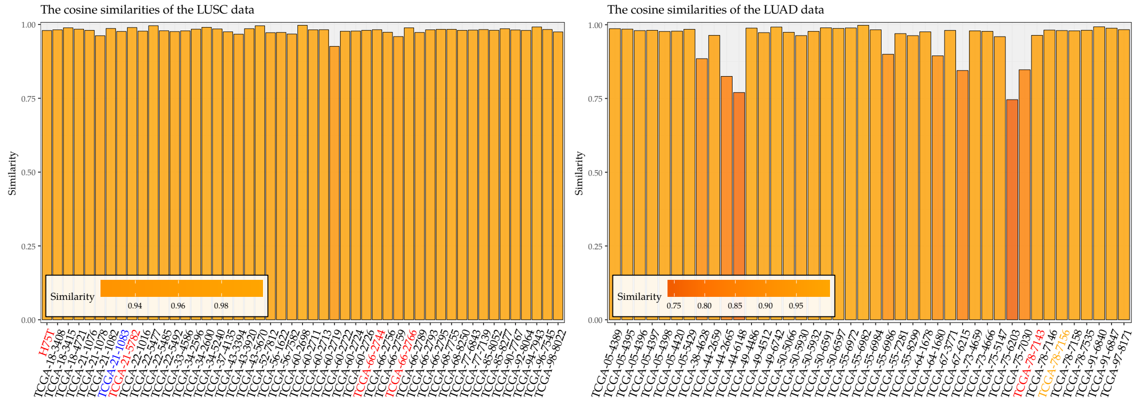


**Suppl.Fig. 7:** *Cosine similarities between the original and reconstructed mutational alphabets. Left: LUSC, Right: LUAD*

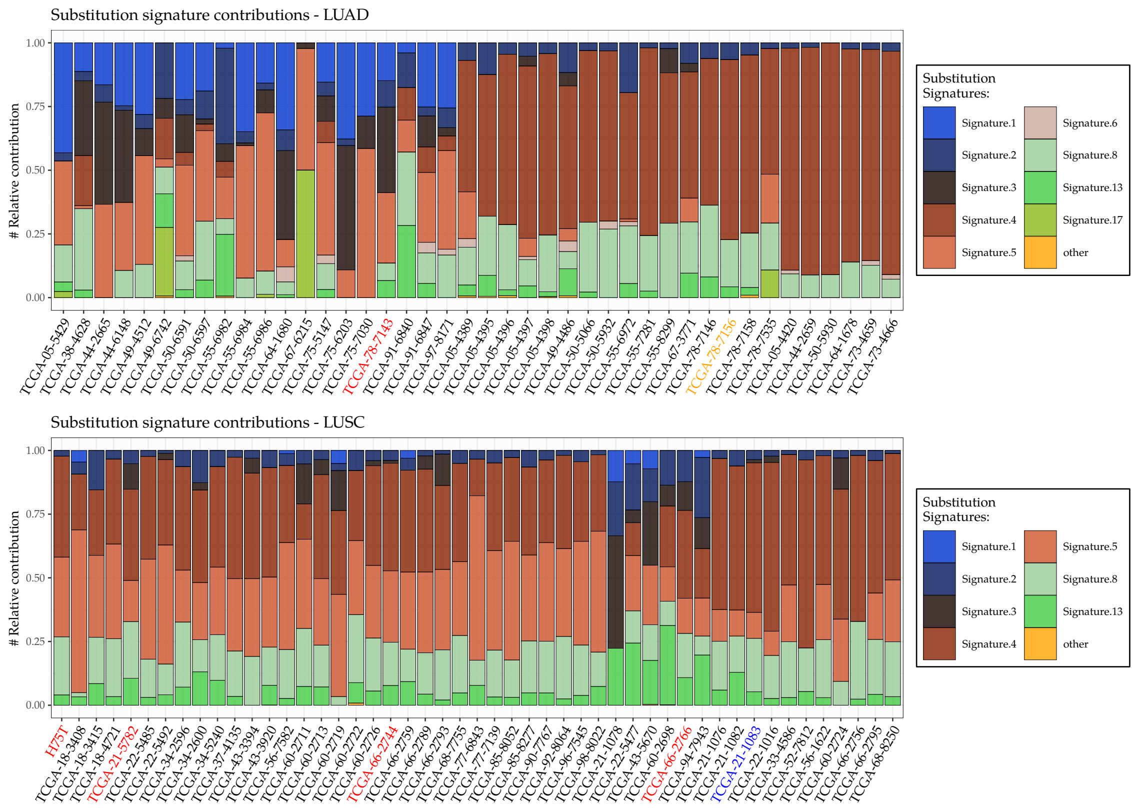


**Suppl.Fig. 8:** *Somatic signature composition of the LUAD and LUSC whole genomes. Top panel: LUAD, Bottom panel: LUSC.*

In general, deletions were classified into three sets: (1) complete repetitions; when the complete deleted sequence is repeated after the deletion in the reference genome, (2) microhomologies; when only the first $n$ nucleotides of the deleted sequence is repeated after the deletion and (3) unique deletions, when the sequence following the deletion has no resemblance to the deleted series of nucleotides. However, since the repetition of the first 1-2 nucleotides could occur by pure chance (With 0.25 and 0.0625 probabilities respectively - assuming that all 4 nucleotides can occur with the same probability), when investigating the effects of the MMEJ/SSA pathway, it is considered a good practice to work with the $n \geq 3$ microhomologies only. Suppl.Fig. **??** provides a summary of this analysis.

#### 4.3 Rearrangement Signatures

Structural Variants had been called using BRASS (v5.4.1). In the analysis only those variants were considered, whose reads could be denovo-assembled by velvet, and at least 6 read-pairs had supported them.

The resulting structural variants then were mapped to the rearrangement-signature alphabet [**?**], and a non-negative least-squares strategy was executed to extract their signature composition. Similarly to the substitution signatures, the similarity between the reconstructed and original spectra was quantified using the cosine between their two 32-dimensional vectors. Since the currently available list of rearrangement signatures is based on breast cancer whole genomes, it isn't surprising, that the cosine similarities of the reconstructions were generally low, especially in the LUAD cohort: mean(cosSim(LUAD)) = $0.77 \pm 0.25$, mean(cosSim(LUSC)) = $0.89 \pm 0.07$. A summary of the structural variants and their rearrangement signature composition is displayed in Suppl.Fig. **??**.

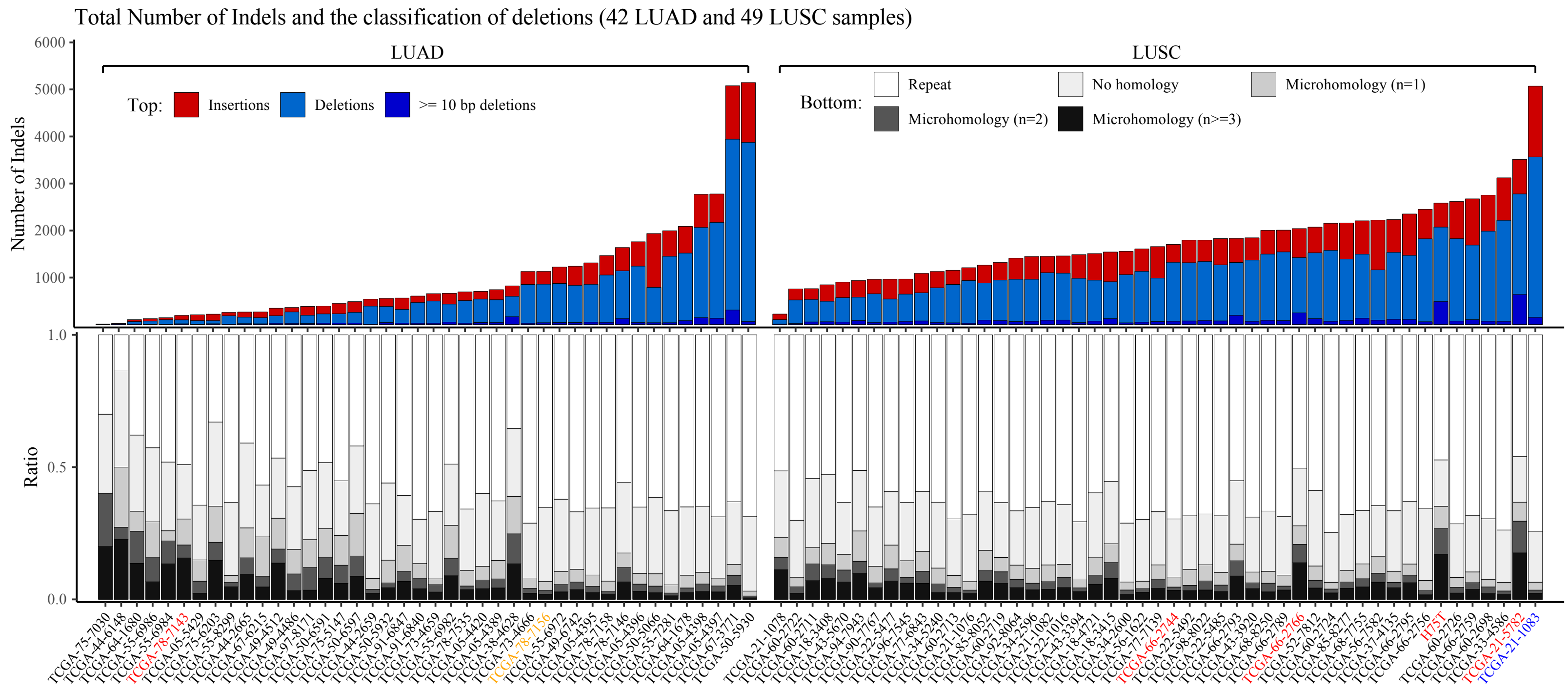


**Suppl.Fig. 9:** *Top panel: Total number of indels in the LUAD and LUSC whole genomes. The horizontal axis contains the absolute number of hard-filtration-passing indels, the bars are colored according to their insertion/deletion content. Deletions are futher divided into <10 and ≥ 10 bp deletions.*

*Bottom panel: Relative constituents of deletions according to the deletion-classification scheme. The three major categories are repeats, microhomologies and deletions without homologous recurrences. Microhomologies are further divided into n=1 bp, n=2 bp, and n≥3 bp variants.*

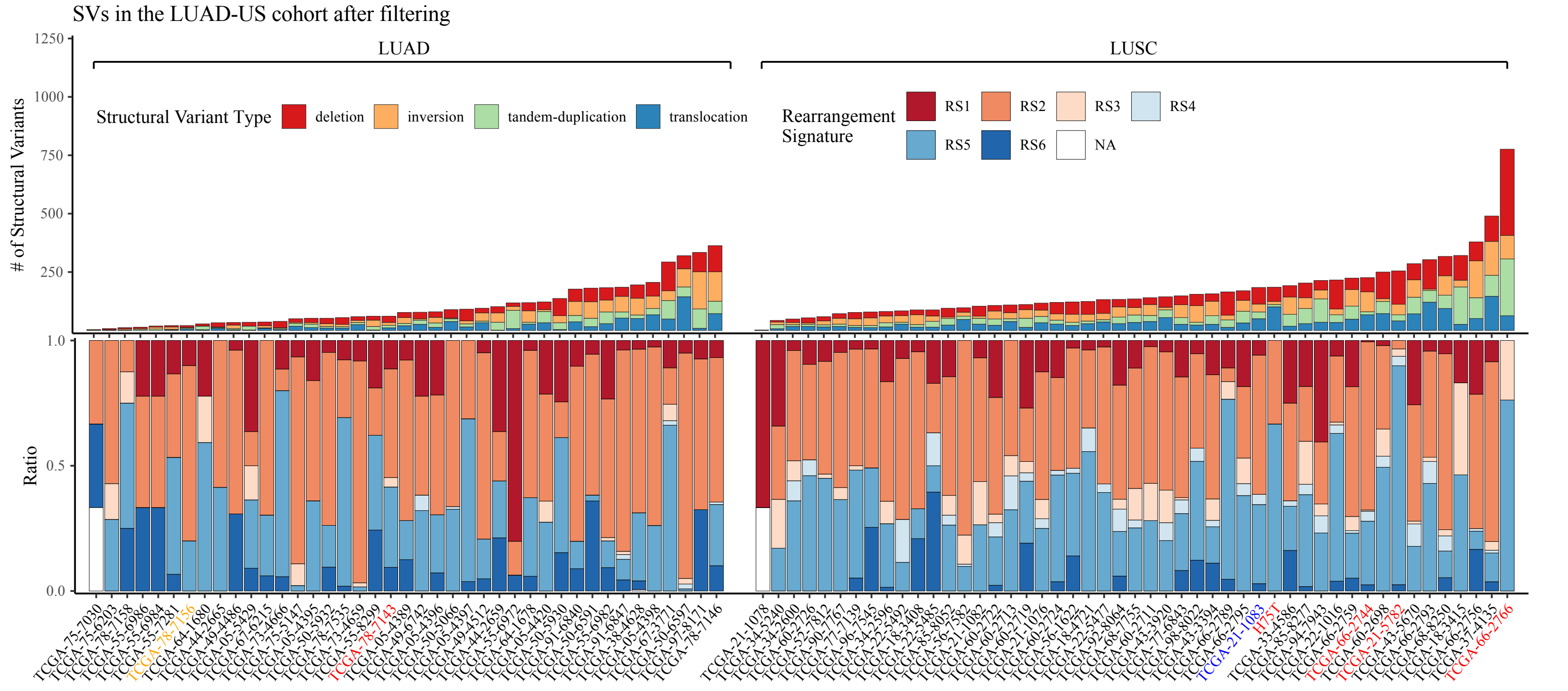


**Suppl.Fig. 10:** *Top panel: Hard-filter-passing structural variants present in the LUAD and LUSC chohorts. The vertical axis shows the total number of structural variants in each sample, on the horizontal axis samples are sorted according to this number. The colors of the bars indicate the relative ratio of each type of structural variants.*

*Bottom panel: Rearrangement signatures in the two lung cancer cohorts. The bars only show the relative compositions, the order of the samples follow the order of the structural variant (top) plot.*

##### 4.4 Genomic scar scores

We have used the `sequenza` R-package [?] to estimate the copy number profiles of the non-small cell lung cancer cohorts. Sequenza can utilize the whole context of whole exome and whole genome sequences, and as such requires the original normal and tumor binary alignment files (BAMs) along with the reference fasta (grch37 in the whole genome and grch38 in the whole exome cases) file that was used for the alignment to do its analysis. When it is done, it provides an estimated allele specific copy number profile for the sample, with the segments corresponding to the parental alleles stored in a data frame. The three genomic scar scores had been calculated from these data frames [?, ?, ?]. The scores had been determined using the scarHRD [?] R package.

##### 4.5 HRDetect scores

Since the whole genome cohort was too small, and the whole exome cohort didn't have enough BRCA mutants, we could not train a new logistic regression model. Instead, we used the original, breast-cancer-specific weights. For the whole genomes:

$$\begin{aligned}
\text{intercept} &= -3.3642 \\
\text{Signature.8} &= 0.09062 \\
\text{HRD-LOH} &= 0.6666 \\
\text{RS5} &= 0.8467 \\
\text{RS3} &= 1.1532 \\
\text{Signature.3} &= 1.6114 \\
\text{mhm.del.ratio} &= 2.3977
\end{aligned}$$

For the whole exomes, we have used a different model, trained on 560 artificially derived (from whole genomes) breast cancer whole exomes [?]:

$$\begin{aligned}
\text{intercept} &= -2.6192939 \\
\text{Signature.17} &= 0.067098 \\
\text{Signature.20} &= 0.09409 \\
\text{Signature.26} &= 0.16166 \\
\text{Signature.6} &= 0.310146 \\
\text{Signature.18} &= 0.31205 \\
\text{mhm.del.ratio} &= 0.314225 \\
\text{Signature.8} &= 0.61474 \\
\text{Signature.13} &= 0.83017 \\
\text{Signature.3} &= 2.00757 \\
\text{HRD-LOH} &= 2.3865
\end{aligned}$$

Before they were seeded to the logistic models, sample attributes had been standardized and log-transformed, just as they were in the original paper [?], however this could had been done in two ways. The first, which is reported in the main article is when the standardization step contains the lung cohorts (LUAD or LUSC separately) only. Since the distributions of the HRD-related attributes is most likely different than the distributions present among breast cancer samples, this form of standardization makes more biological sense.

However, we argued, that it is worth to check what would be the HRDetect status of these samples, if we would treat them as breast cancers. In order to check this, the two lung cohorts had been appended to the 560 breast WGS dataset, and the standardization was executed on the resulting 602 (breast + LUAD) and 608 (breast + LUSC) specimens. The resulting distribution of "breast-standardized" HRDetect scores is displayed on Suppl.Fig. ??. In this scenario only a single LUSC sample (TCGA-66-2766) exceeds the 0.7 HRDetect score threshold.

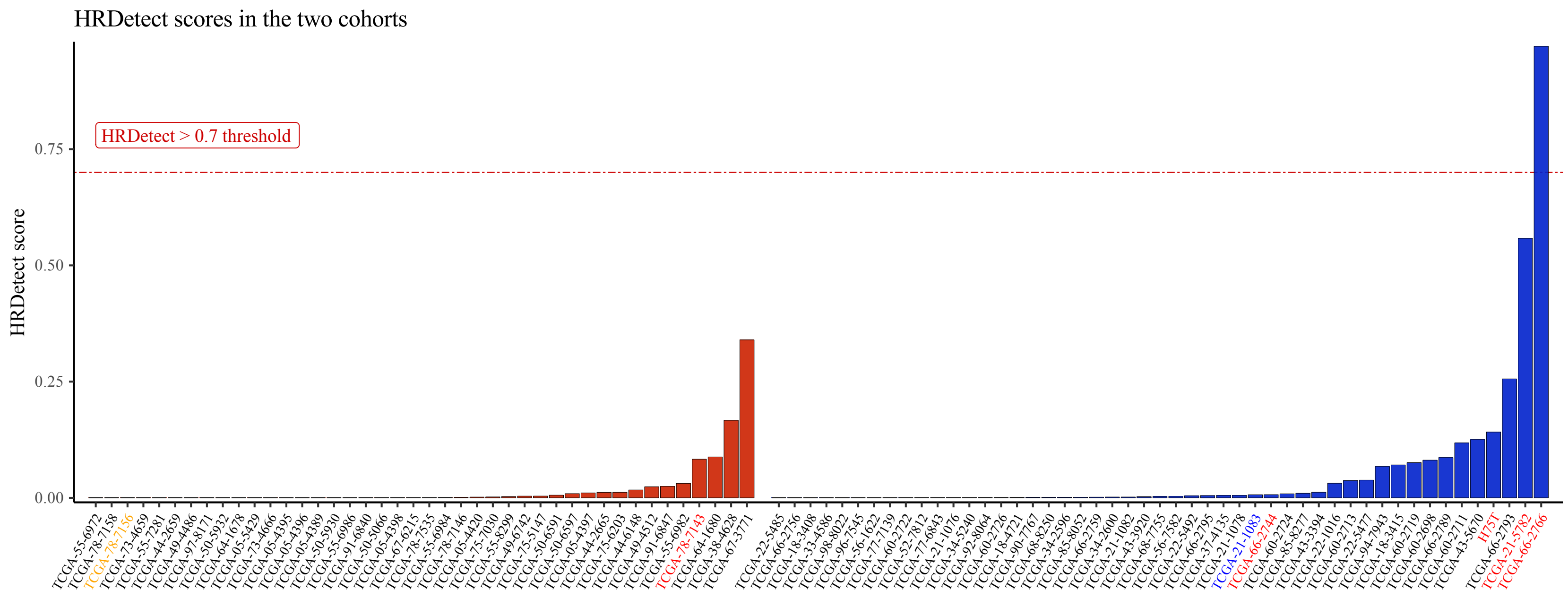


**Suppl.Fig. 11:** *Breast cancer standardized HRDetect scores of the LUAD and LUSC whole genomes.*

### 5 HRD-related genomic features extracted from the whole exomes

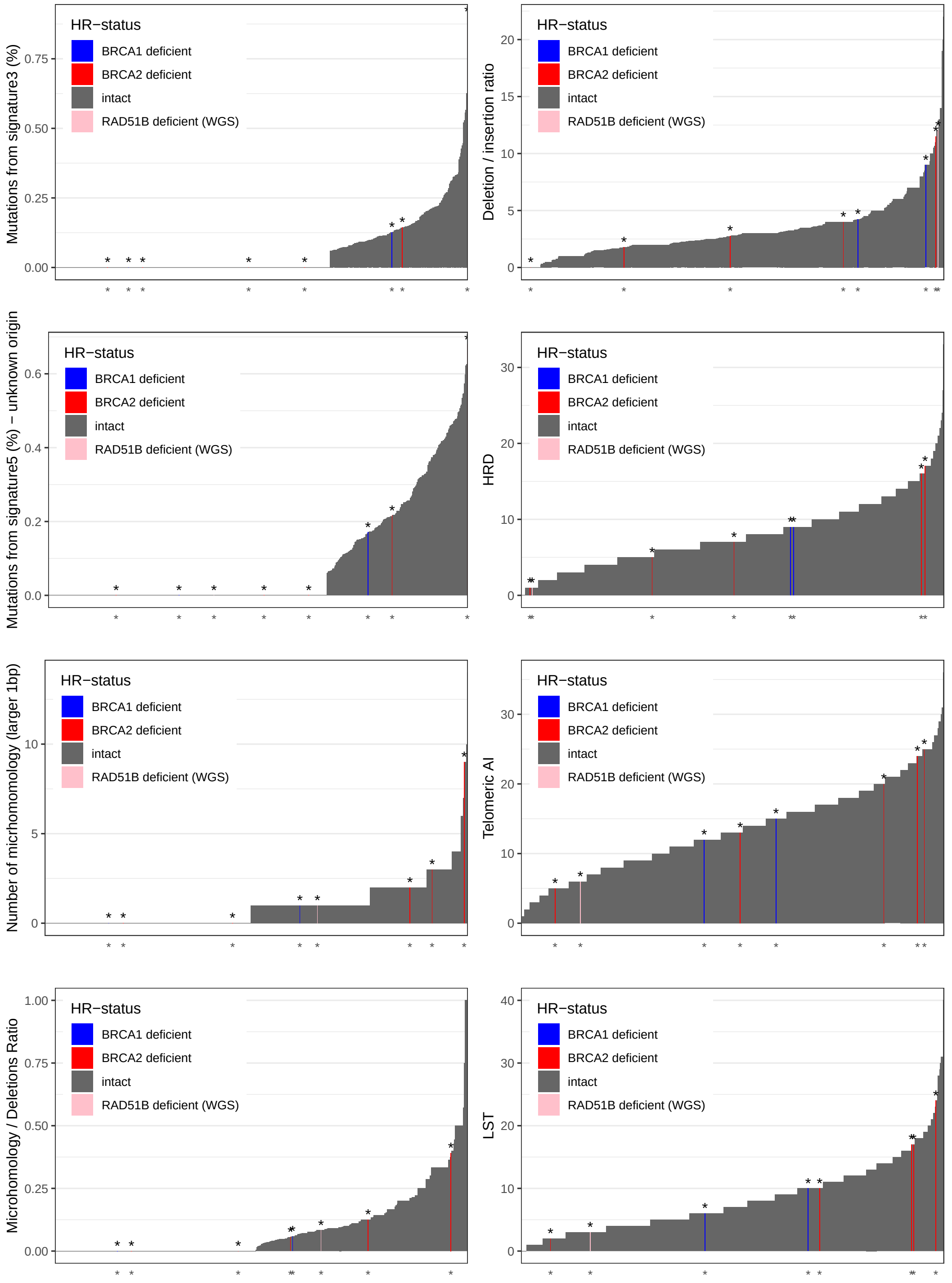


**Suppl.Fig. 12:** *Distribution of genomic scar scores (HRD-LOH, Telomeric Allelic Imbalance, Large-scale Transitions), homologous-recombination deficiency related mutational signatures (Signature 3, 5), number of micrhomomology-mediated deletions, microhomology / deletions ratio, deletion / insertion ratio and BRCA1/2-status in whole exome sequenced lung adenocarcinoma samples (n=553).*

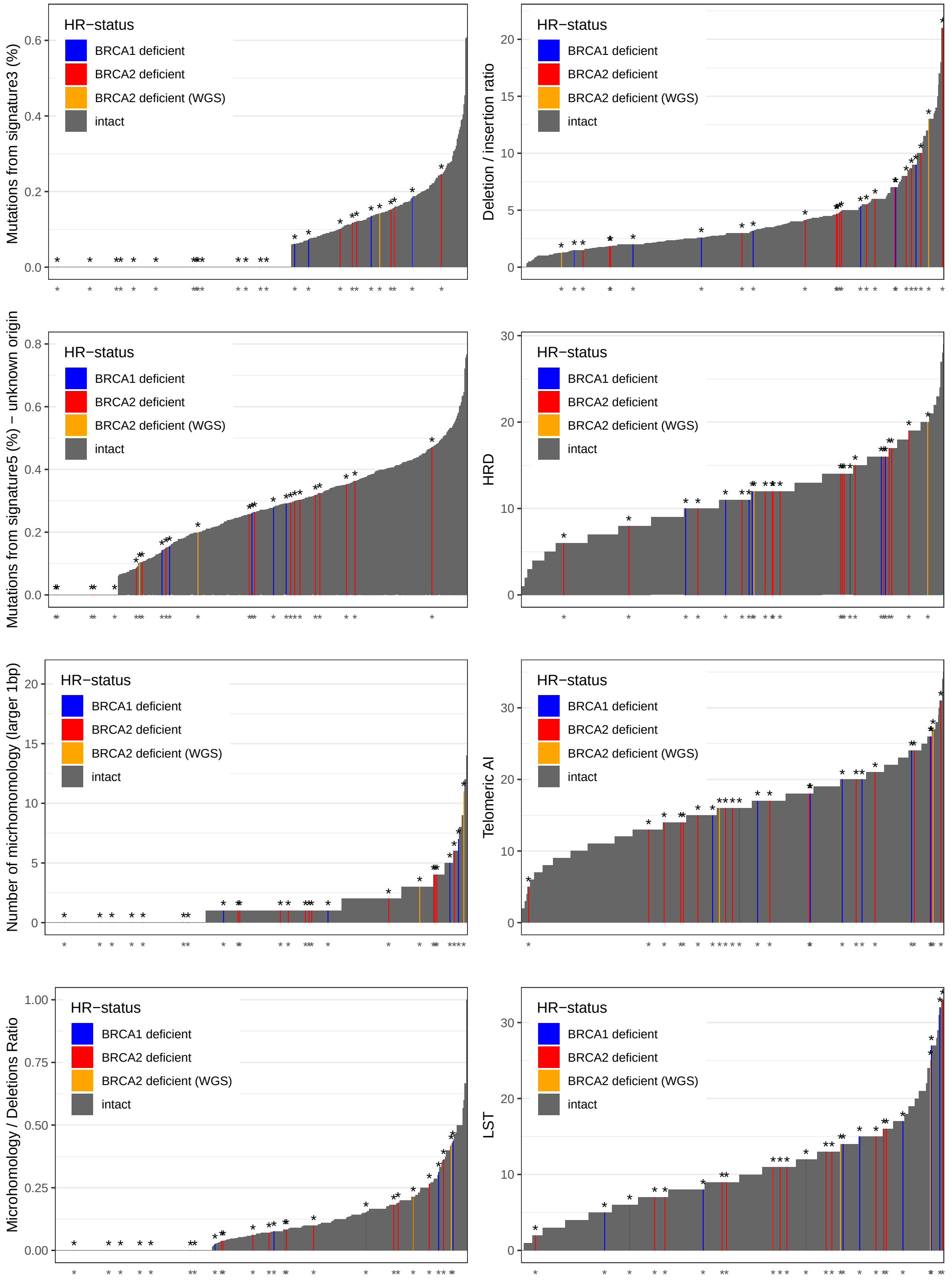


**Suppl.Fig. 13:** *Distribution of genomic scar scores (HRD-LOH, Telomeric Allelic Imbalance, Large-scale Transitions), homologous-recombination deficiency related mutational signatures (Signature 3, 5), number of micrhomomology-mediated deletions, microhomology / deletions ratio, deletion / insertion ratio and BRCA1/2-status in whole exome sequenced lung squamous carcinoma samples (n=489).*

### 6 Correlations between the WES and WGS HRD-predictors

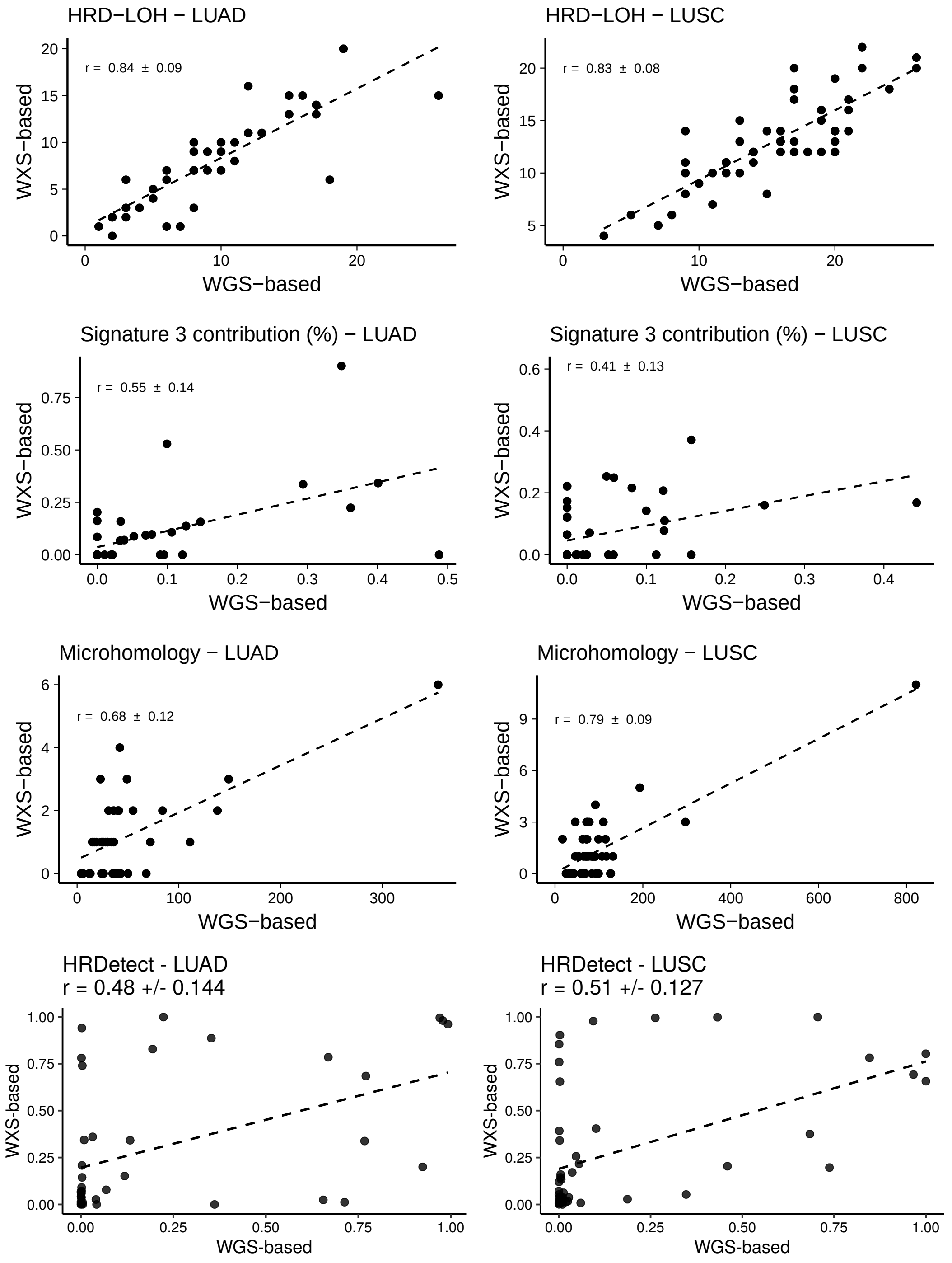


**Suppl.Fig. 14:** *Correlation of the three main components (number of HRD-LOH events, number of microhomology-mediated deletions, and contribution of Signature 3 to the mutational profile) of HRDetect between paired whole exome and whole genome sequenced samples*

### 7 HRDetect-scores in TCGA WES cohorts

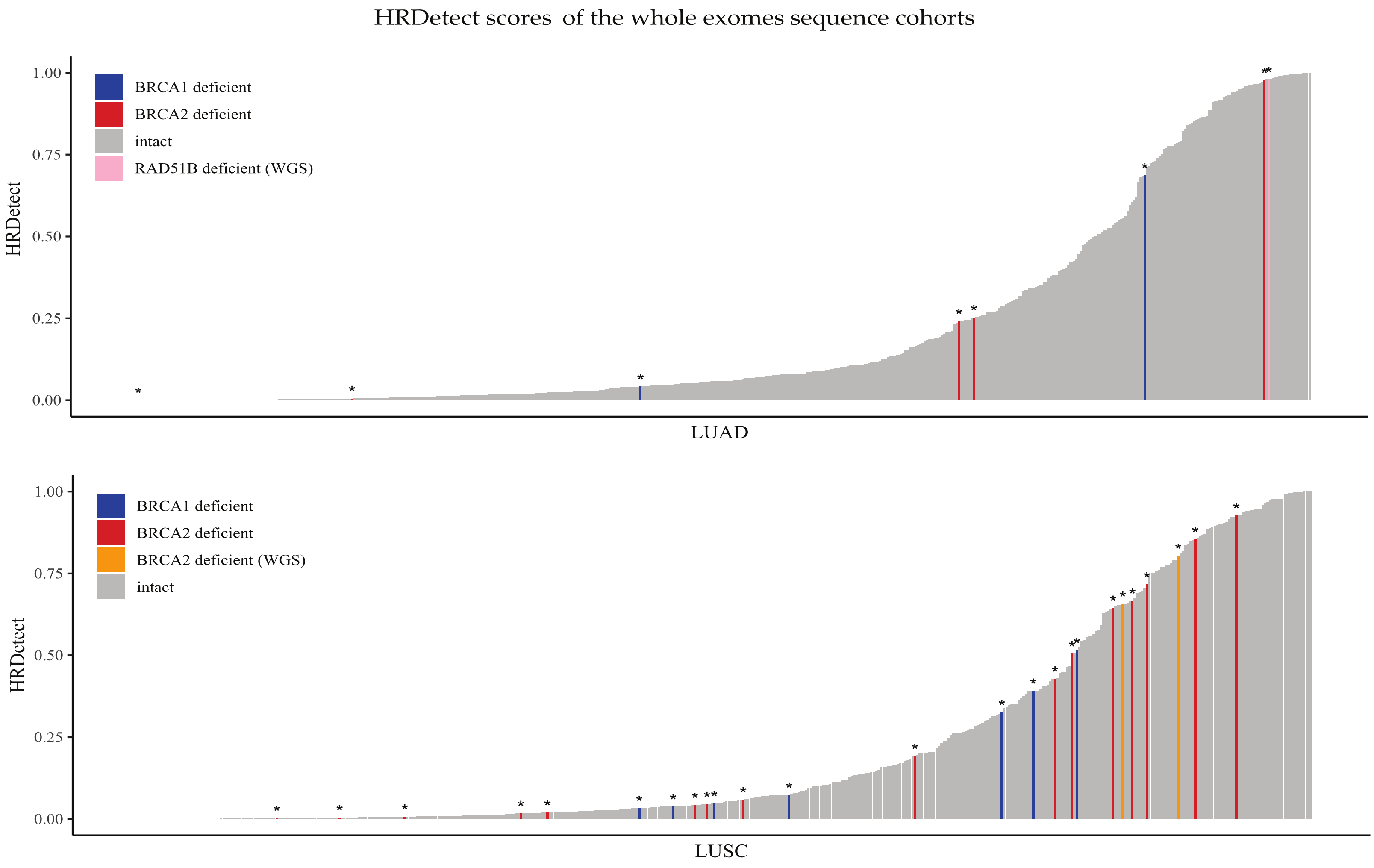


**Suppl.Fig. 15:** *Breast cancer standardized HRDetect scores of the LUAD and LUSC whole exome (TCGA).*

##### 7.1 Comparison of WES and WGS HRDetect-scores

The two LUSC patients showing signs of HR-deficiency based on whole genome sequencing, also had high HRDetect values based on whole exome sequencing analysis (TCGA-21-5782: 0.80, TCGA-66-2766: 0.66). Since the HR deficiency status of these two cases are supported by WGS data, we used the lower HRDetect value of these two cases as a putative threshold for HR deficiency in the WES characterized LUSC cohort. In the LUSC WES cohort 16% of the patients had higher than 0.66 HRDetect scores (Figure 3B), while in the case of the LUAD cohort (Figure 3A) 3.8% of the patients had at least as high HRDetect score as the RAD51B-mutated sample (TCGA-64-1680).

#### 8 Survival analysis

Higher WXS-based HRDetect-score was not associated with better progression free survival (PFS) or overall survival (OS) in LUAD and LUSC patients in the TCGA dataset. There was also no significant difference among the subset of patients who received platinum treatment.

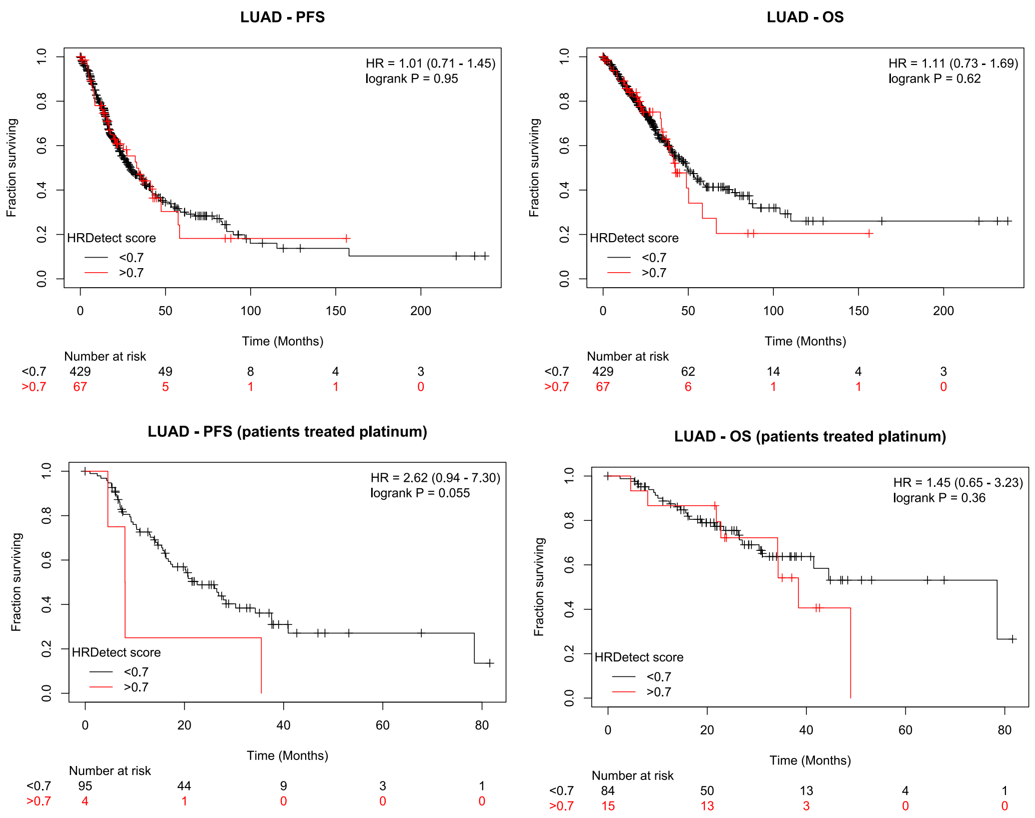


**Suppl.Fig. 16:** *Survival curves - LUAD*

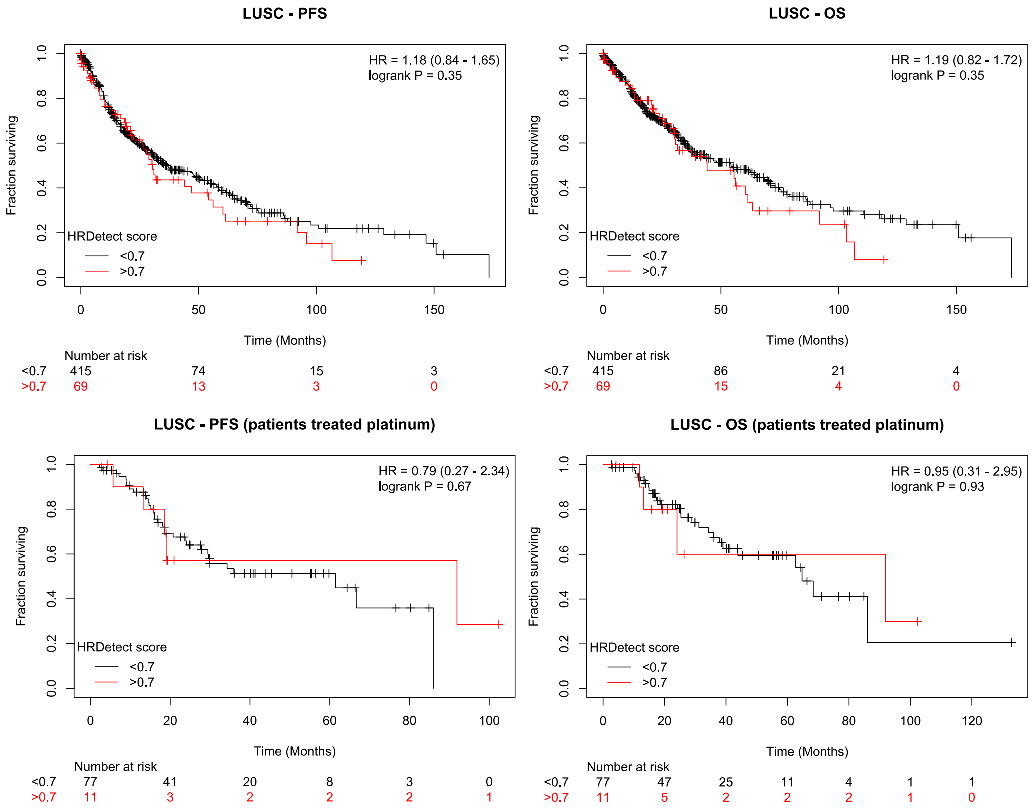


**Suppl.Fig. 17:** *Survival curves - LUSC*

### 9 HRDetect-score in cell lines

The simplified HRDetect model for cell lines using only single-nucleotide mutational signatures and the proportion of microhomology-mediated deletions were retrained on the 560 breast cancer artificial WES data.

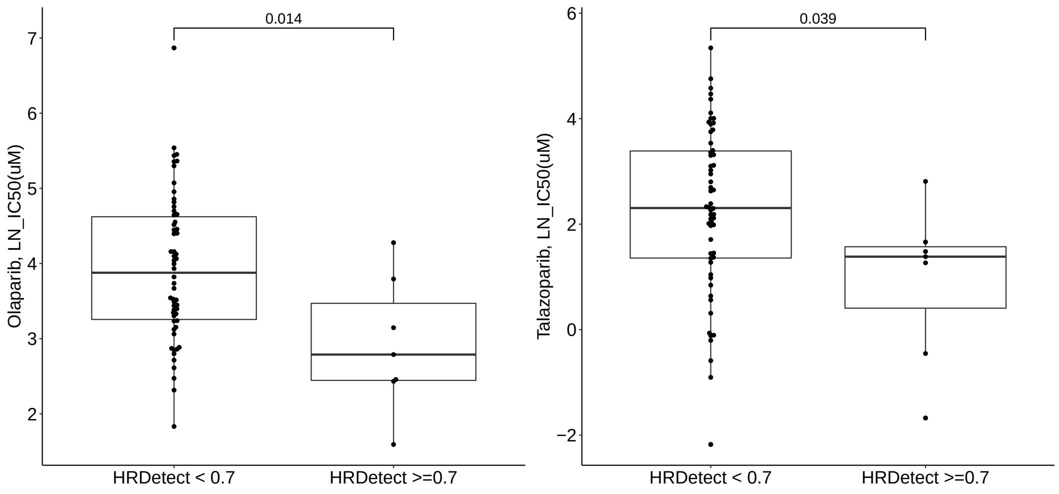


**Suppl.Fig. 18:** *Lung cancer cell lines with larger than 0.70 HRDetect scores showed significantly (p<0.05) higher sensitivity to olaparib and talazoparib based on CCLE [**?**] and GDSC [**?**] data. x-axis: Natural log of the fitted IC50(uM)*
